# Gills, growth, and activity across fishes

**DOI:** 10.1101/2022.09.24.509310

**Authors:** Jennifer S. Bigman, Nicholas C. Wegner, Nicholas K. Dulvy

**Author notes:** Alaska Fisheries Science Center, NOAA, 7600 Sand Point Way NE, Seattle, WA 98115.

## Abstract

Life history theory argues that an organism’s maximum size and its corresponding growth rate have evolved to maximize lifetime reproductive output. The Gill Oxygen Limitation Theory suggests that in aquatic organisms, maximum size is instead constrained by the surface area of the gills, the primary site of gas exchange with the environment. A central prediction of this theory is a tight relationship among maximum size, growth, and gill surface area. Yet since this idea was first tested in the early 1980s, data availability has increased and analytical methods have advanced considerably. Here, we revisit this relationship with new data and a novel phylogenetic Bayesian multilevel modeling framework that allows us to understand how individual variation in gill surface area confers relationships of maximum size, growth, and gills across species. Specifically, we bring gill surface area into an allometric context and examine whether the gill surface area for a given body size (intercept) and the rate at which gill surface changes with size (slope), for a given species, explains growth performance -- an index integrating the life history tradeoff between growth and maximum size -- across fish species. Additionally, we assess whether variation in von Bertalanffy growth coefficients across species can be explained by gill surface area. Finally, we explore whether additional factors – here, activity and evolutionary history -- explain variation in maximum size and growth across species. Overall, we find that although a positive relationship exists among maximum size, growth, and gill surface area across fishes, it is weak. Additionally, gill surface area does not explain much variation in growth coefficients across species, especially for those that reach the same maximum size. However, we find that the activity level of a fish explains more variation in maximum size and growth across species compared to gill surface area. Our results support the idea that in fishes, growth and maximum size are not simply related to gill surface area, and that other covariates—both tractable (e.g., activity, metabolic rate, temperature) and less tractable (e.g., predation risk, resource availability, and variation)—appear to explain more variation in life history traits across species.

## Introduction

Formalized as life history theory, decades of work have revealed that body size, and other life history traits related to growth, survival, and reproduction, are optimized by natural selection to maximize fitness (typically measured by reproductive output in fishes; Beverton & Holt 1959; Stearns 1992, Hutchings 2002). Maximizing fitness results in tradeoffs between traits (such as growth and reproduction) as competing processes draw from the same finite pool of internal resources (e.g., time, energy; Roff 1984, Stearns 1989, Reynolds 2003). One of the classic tradeoffs between life history traits is the inverse relationship observed between the maximum size of a species and its change in body size over time, or growth (Beverton & Holt 1959, Reynolds *et al*. 2001). This tradeoff suggests that an individual (or species) generally grows faster to a smaller asymptotic (final) size or grows more slowly to a larger asymptotic size (Beverton & Holt 1959). With respect to maximum size and growth tradeoffs, life history theory predicts that under high mortality (e.g., in an unstable environment or under high predation risk) fitness would be maximized through a faster life history strategy, one that results in a higher reproductive output earlier in life, which would select for a smaller maximum size, faster growth, and earlier maturity (Stearns 1976, Roff 1984; Reznick *et al*. 1996). On the other hand, under low mortality (e.g., a stable environment or one with lower predation risk) fitness would be maximized through a slower life history strategy by waiting to reproduce until an organism reaches a larger size (as reproductive output increases with increasing size; Bjørkvoll *et al*. 2012, Barneche *et al*. 2018). This would select for a larger maximum size, slower growth, and later maturity (Stearns 1976, Roff 1984). While life history theory and its predictions have been widely supported by both theoretical and empirical research over the last 70 years, recent work on the effect of oxygen (i.e., the balance of supply and demand) and temperature on body size and growth, especially for fishes, has inspired the proposal of a new mechanistic theory that shapes body size and growth (Pauly 2010, Forster *et al*. 2012, Cheung *et al*. 2013).

The Gill Oxygen Limitation Theory proposes that the maximum size of aquatic, water-breathing organisms is mechanistically constrained by oxygen supply at the gills (Pauly 1981, 2010, 2021). The central tenet of this theory is that the oxygen supply acquired over the surface area of the gills—which is (to a first approximation) a two-dimensional surface—cannot keep pace with the demand from a continually increasing three-dimensional volume (body mass). The proposed consequence of this mismatch in geometry is that the ontogenetic slope of the relationship of gill surface area and body mass will always be less than one (i.e., hypoallometric). This means that the ratio of gill surface area to body mass (i.e., mass-specific gill surface area) will decrease with increasing body mass. Thus, when the supply of oxygen diffused over the ‘diminishing’ gill surface area cannot match the demand from the growing body, the organism will stop growing and its maximum size will be reached (Pauly 2010, 2021). Indeed, the Gill Oxygen Limitation Theory is rooted in the von Bertalanffy growth model such that growth is a function of anabolism (synthesis of material) and catabolism (breaking down of material). The von

Bertalanffy growth model is based on the idea that growth occurs when anabolism is greater than catabolism and growth stops when anabolism equals catabolism. The Gill Oxygen Limitation Theory argues that because anabolism requires oxygen, and catabolism does not, growth can be thought of as a function of anabolism, which ultimately, is driven by the amount of oxygen that a fish or other water-breathing organism can diffuse over the surface area of the gills (Pauly 1981, 2021). Thus, this theory suggests that the mechanism underlying the process of growth ceasing due to anabolism equaling catabolism is a function of oxygen supply via the gills (Pauly 1981, 2021).

Because of this connection to the von Bertalanffy growth function, a central prediction of the Gill Oxygen Limitation Theory is that a tight relationship exists among maximum size, growth, and gill surface area (Pauly 1981, Pauly 2010). This relationship was first tested over 40 years ago by examining whether growth performance (an index that integrates the tradeoff between the von Bertalanffy growth coefficient and asymptotic size) explained variation in a proxy of gill surface area (gill area index; Pauly 1981). Further, it was suggested that the large amount of variation in von Bertalanffy growth coefficients both within and across species was related to gill surface area, such that an individual or a species can only grow fast to its asymptotic size if it has a larger than expected gill surface area for its body size (Pauly 1981, 2010). Although only a weak positive relationship between gill area index and growth performance existed in the 42 fish species originally examined by Pauly (1981), this relationship underpinned the idea that gill surface area constrains growth and maximum (or asymptotic) size in fishes and has been used, in part, to predict future changes in fish maximum size associated with ocean warming (Pauly 1981, 2010, 2021, Cheung *et al*. 2013, Cheung & Pauly 2016). Indeed, half of the predicted 14 – 24% decline in maximum size for individual fish species (over generations) due to projected temperature increases through 2050 has been suggested to be mechanistically linked to oxygen limitation, or the mismatch between oxygen supply (gill surface area) and demand (metabolic rate; Pauly 1981, 2010, Cheung *et al*. 2013). This ‘shrinking fishes’ finding awakened renewed interest in the Gill Oxygen Limitation Theory (Bigman et al. 2021, Lefevre et al. 2021, Pauly 2021, Roche et al. 2022).

Much of the interest in the Gill Oxygen Limitation Theory is centered on the debate surrounding the mechanistic underpinnings of oxygen limitation, particularly with respect to gill surface area (Lefevre *et al*. 2017, 2018, Marshall & White 2019). The classical physiological view is that the surface area of respiratory organs evolve to provide the capacity needed to meet an organism’s requirements, instead of (aerobic) metabolic rate being driven by, and ultimately, limited by the surface area of the gills (Lefevre *et al*. 2017, 2018, Marshall & White 2019). Relatedly, physiologists have noted that the surface area of gills are folded surfaces and thus are not under the same strict geometric constraints as seen in spherical objects (i.e., the scaling of gill surface area and body mass can and do deviate from theoretical surface area-to-volume ratios; Wegner, 2011, Lefevre *et al*. 2017, 2021). On a more technical note, the proxy for gill surface area originally used to test whether a relationship between growth performance and gill surface area existed – gill area index -- does not capture the known (large) variability in gill surface area within and across species and is biased by the sizes at which gills were measured (see Bigman et al. *in review*). Notwithstanding these criticisms, the Gill Oxygen Limitation Theory has potentially far-reaching consequences if empirically supported. In addition to the idea that oxygen limitation and gill surface area may be behind the observed declines in maximum size in response to increasing temperature (temperature-size rule/Bergmann’s rule/James’ rule), mounting evidence from broad, cross-species studies suggests that oxygen limitation may also shape species’ geographic distributions and underlie the mass- and temperature-dependence of metabolic rate (Forster *et al*. 2012, Deutsch *et al*. 2020, Rubalcaba *et al*. 2020, Bigman *et al*. 2021, Clarke et al. 2021, English et al. 2022, Essington et al. 2022).

To understand whether oxygen limitation mediated by gill surface area is indeed occurring, and affecting growth and maximum size, predictions generated by the Gill Oxygen Limitation Theory must be tested. Yet, few predictions have been tested to date, including the generality of the relationship among maximum size, growth, and gill surface area. A wealth of gill surface area and life history data have been published over the last 40 years, with many species possessing raw gill surface area data—or measures for multiple individuals of the same species. Additionally, there has been an advancement of statistical techniques that can incorporate additional salient factors, such as evolutionary history among species, and allow us to link individual variation to patterns across species. To that end, we revisit the relationship of maximum size, growth, and gill surface area in fishes by leveraging available gill surface area data and developing a novel phylogenetic Bayesian multilevel model with the flexibility to scale up individual variation to assess patterns across species, as well as include salient covariates. Specifically, we ask three questions: (1) do species with faster growth coefficients for their body size have larger gills, (2) do estimates of gill surface area that incorporate its size-dependency (the slope and intercept of the ontogenetic allometry) relate to growth performance across fishes, and (3) does activity level better characterize the variation in growth performance across species compared to gill surface area?

## Methods

### Additional data collection and sources

We compiled a dataset of species-specific gill surface area estimates with their associated body masses at measurement (“measurement body mass”) and von Bertalanffy growth parameters for both teleosts and elasmobranchs. An initial dataset was collated for those fish species with estimates of both gill surface area *and* available growth parameters in Fishbase (Froese and Pauly 2019). This initial dataset was then supplemented with published gill surface area data from other sources (if a given species also had available growth parameters). These other sources of gill surface area data were: Gray (1954), Hughes & Morgan (1973), De Jager and Dekkers (1975), and Palzenberger & Pohla (1992), Bigman *et al*. (2021), and references therein. We additionally limited our dataset to species that have a resolved position on a phylogenetic tree to allow for including a random effect of evolutionary history in our models.

### Gill surface area data

Gill surface area estimates (cm^2^ or mm^2^) and measurement body mass were extracted from the original study in which they were reported. Only those species with raw gill surface area data (i.e., estimates for multiple individuals of a species, each with its own measurement body mass) were included in our dataset. If more than one study reported raw data for a number of individuals for a given species, we combined both datasets (this was only the case for three species: *Alopias vulpinus* Common Thresher Shark, *Carcharhinus plumbeus* Sandbar Shark, *Isurus oxyrinchus* Shortfin Mako). Any gill surface area estimate that was not directly measured (e.g., predicted from assumed geometric relationships) was not included in this study (for further discussion see Satora and Wegner 2012). All gill surface area and body mass estimates were converted to cm^2^ if not already in this unit and log_10_ -transformed prior to analyses.

### Life history data

Using the ‘rfishbase’ package for Fishbase, we extracted all observations of von Bertalanffy growth function parameters for each species in our dataset including the growth coefficient, *k* (year^-1^) and asymptotic length (the mean length the individuals in a population would reach if they were to grow indefinitely), L (cm; Froese and Pauly 2019, Boettiger *et al*. 2012). If the type of length (i.e., total length, TL, fork length, FL, etc.) was not specified for an observation of l<)(), then that observation (both *k* and l<)() could not be used and was removed from the dataset. If growth data were not available in Fishbase for a species, the primary literature was searched for published age and growth data. For most species, the asymptotic size was reported as L and not and thus was estimated for each observation using length-weight regressions matched by length type and sex downloaded from Fishbase using the ‘rfishbase’ package (Boettiger *et al*. 2012). For one species (*Torpedo marmorata*), growth parameters were not available in Fishbase but were found in the literature. For four species, length-weight coefficients for the same length type as was used to estimate growth parameters were not available (i.e., the growth coefficient was estimated using fork length but no length-weight regression for fork length to weight was found [and no conversion from another length type was available]), and thus matching type-specific length-weight regressions were collated from the literature. Growth performance was calculated for each observation as log_10_ (*k* * W<)()) following Pauly (1981). For analyses, a mean of growth performance was taken for each species.

### Activity data

To include a standardized, objective, and quantitative metric of activity level in our analyses, we estimated the caudal fin aspect ratio – a morphological correlate of swimming speed and ability – for each species (Palomares & Pauly 1989, Sambilay 1990, Bigman *et al*. 2018). While we recognize that there may be shortcomings of using caudal fin aspect ratio as a metric of activity level (i.e., tail shape may change slightly with growth), this metric offers a straightforward and consistent method to quantitatively index activity level that can be obtained for most species that have an anatomically correct illustration (i.e., from field guides). In contrast, the swimming capacity index used by Pauly (2010) requires species-specific swimming speed data, which does not exist for most species. Caudal fin aspect ratio (*A*) was calculated for each species as *A = h*^2^ */ s*, where *h* is the height and *s* is the surface area of the caudal fin as measured from anatomically correct field guide illustrations from *Sharks of the World* (Ebert *et al*. 2016) for elasmobranch species or FAO field guides for teleost species (Palomares & Pauly 1989, Sambilay 1990, Bigman *et al*. 2018). For some teleost species, FAO images were not available (n = 7) and thus alternative field guides were used. If alternative guides were used, we generated a mean caudal fin aspect ratio from up to four field guide illustrations (if available). Of the 32 species in our dataset, caudal fin aspect ratio could only be estimated for 30 species as two species do not have traditional caudal fin morphology (one eel, *Anguilla anguilla*, and one batoid, *Torpedo marmorata*).

The growth of ‘aquacultured’ species is known to differ from that of wild fishes (due to food ad libitum, reduced predation, and possibly increased aeration of aquaculture ponds; Pauly 2010). Also, fishes that breathe air (either by possessing an air-breathing organ or passive oxygen diffusion through the skin) often have a lower gill surface area for a given body size compared to their non-air-breathing counterparts (Wegner 2011). Thus, we created two subsets of our full dataset to exclude species traditionally used in aquaculture and those capable of air-breathing.

In total, our dataset included 457 observations of raw gill surface area and associated body masses from a total of 32 fish species (both teleosts and elasmobranchs) for which von Bertalanffy growth parameters were available and have a resolved position on a published phylogeny (Table S1). Importantly, these raw data allow us to estimate ontogenetic allometric coefficients (the slope and intercept) of gill surface area that capture the size-dependency of this trait and examine whether gill surface area does indeed relate to growth and maximum size (as opposed to a mean of gill surface area and measurement body mass on an unknown number of individuals over an unknown body size range). However, estimating allometric coefficients must be done with care as too few data points can produce biased coefficients (White and Kearney 2014, Jenkins & Quintana-Ascencio 2020). Indeed, previous work has shown that a threshold of eight individuals is suitable to reliably estimate regression parameters (Jenkins & Quintana-Ascencio 2020). To ensure this was true for our dataset, we simulated regression coefficients with 3 – 100 data points and found that coefficients did not differ substantially past eight samples (see SI, Figure S1). We also reran all models with an additional subset, one in which we filtered according to body size, and only those species that had a body size range of at least an order of magnitude were included (n = 32 species).

All data (and sources) assembled for this study are archived on github (see references section for link to data and code).

### Analyses

#### 1. Do species with faster growth coefficients for their body size have larger gills?

To assess whether gill surface area, in addition to asymptotic size, explained variation in von Bertalanffy growth coefficients across species, we developed a novel Bayesian multilevel modeling framework that included three levels. The first level of the model estimated the ontogenetic allometry of gill surface area and body mass for each species, resulting in a species-specific posterior distribution of the intercept (gill surface area at a given size) and a species-specific posterior distribution of the ontogenetic slope (rate at which gill surface area increased with body mass). The second level of the model then examined whether the gill surface area intercept or ontogenetic slope explained additional variation in the relationship between the von Bertalanffy growth coefficient and asymptotic size across species. To ensure that intercepts were estimated accurately across the broad size range of species included in the dataset, body mass data were centered on the mean value of body mass for all 32 species in the dataset (300 g). Both predictors (the gill surface area intercept or slope and asymptotic size) in the second level of the model were standardized (by z-score) to facilitate comparison and infer the relative importance of a given predictor in explaining variation among growth coefficients. We estimated the correlation and variance inflation factors (VIF) between the gill surface area intercept or slope and asymptotic size for both models to ensure that these traits, as parameterized in our models, were not collinear or correlated. The strength of using such a multilevel modeling approach is that the uncertainty in the species-specific intercepts and ontogenetic slopes estimated in the first level of the model is propagated across levels of the model as each iteration of all levels of the model happens in succession. For more details on our modeling approach, see the SI.

#### 2. Do estimates of gill surface area that incorporate its size-dependency (the slope and intercept of the ontogenetic allometry) relate to growth performance across fishes?

To assess whether growth performance varied with gill surface area intercept or slope, we fit two Bayesian multilevel models. The first level of both models estimated the species-specific posterior distribution of the ontogenetic intercepts and slopes of the relationship of gill surface area and body mass (centered at 300 g, as above). The second level of both models then examined whether the intercept or the slope of the ontogenetic allometry explained variation in growth performance across species. In both models, the species-specific slopes and intercepts were standardized using the z-score transformation for input in the second level of the model, which facilitated model convergence and parameter estimation.

#### 3. Does activity level better characterize the variation in growth performance across species compared to gill surface area?

To assess whether activity level explained more variation in growth performance compared to gill surface area, we fit two multilevel Bayesian models which allowed for inferring the relative importance of each predictor (caudal fin aspect ratio vs. gill surface area intercept or slope). The first level of both models estimated the species-specific posterior distribution of the ontogenetic intercepts and slopes of the relationship of gill surface area and centered body mass (at 300 g, as above). The second level of the two models examined whether caudal fin aspect ratio or the ontogenetic gill surface area intercept (model 1) or the ontogenetic gill surface area slope (model 2) explained more variation in growth performance. All predictors were standardized, and we estimated the correlation and variance inflation factors (VIF) for both models to ensure that activity level and gill surface area, as parameterized in our models, were not collinear or correlated.

We ran all models in all three questions above with and without a random effect of phylogeny to ensure our results were not biased due to species’ sharing various parts of evolutionary trajectories (e.g., see Felsenstein 1985; Freckleton 2009; Harmon 2018). To do so, we constructed a new supertree with species from our dataset using two published phylogenies -- one for teleosts (Rabosky *et al*. 2018, Chang *et al*. 2019) and one for chondrichthyans (Stein *et al*. 2018). Models with and without a random effect of phylogeny yielded almost identical results (Tables 1-3).

**Table 1.**
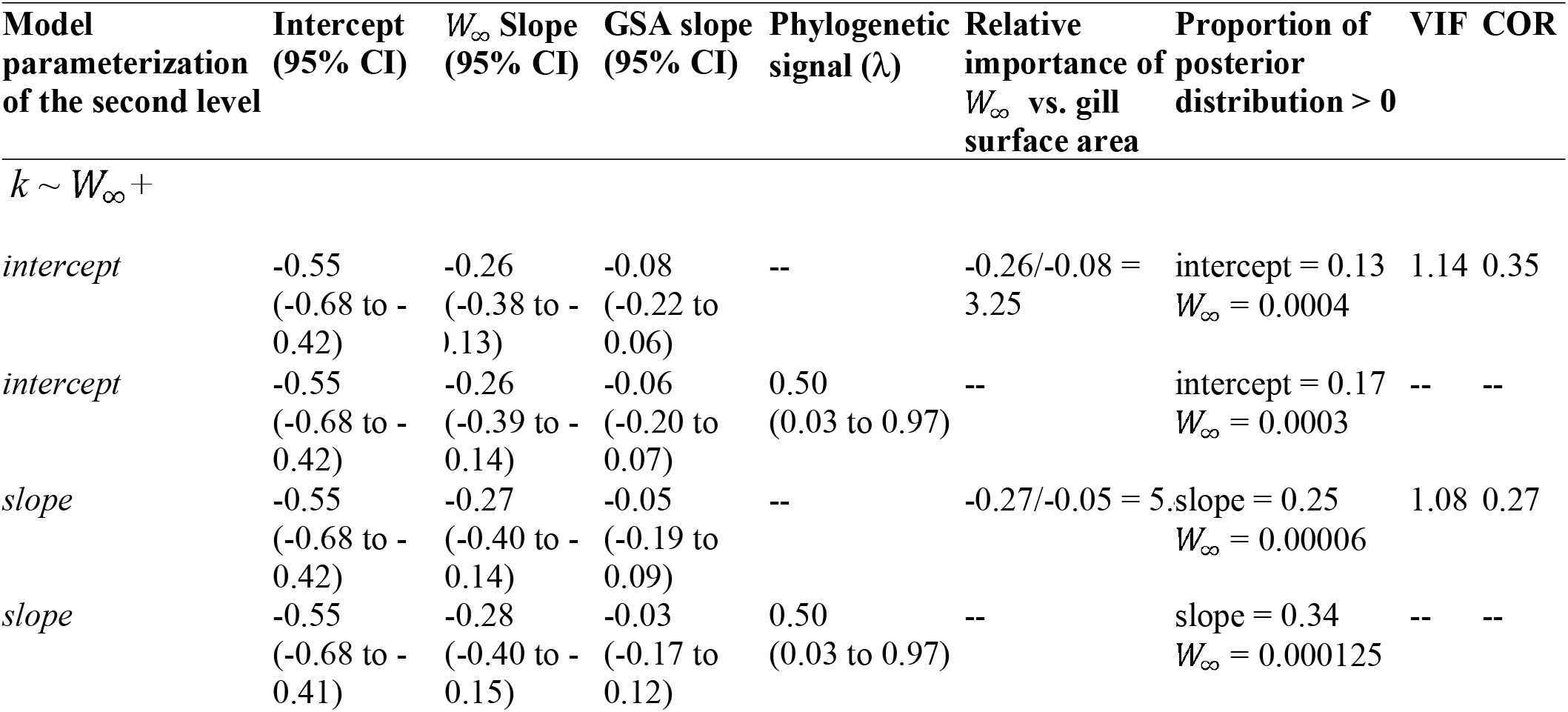
Comparison of coefficients and 95% Bayesian Credible Intervals (BCI) for the relationship of the growth coefficients (*k*), asymptotic size (WOO), and gill surface area, as measured by the ontogenetic intercept or slope. All models were estimated using a Bayesian multilevel modeling framework in Stan using the package *rstan* in R v.4.0.2. All predictors in the second level of the model were standardized and thus the effect sizes for the slopes are relative to each other (see text and SI). GSA = gill surface area, VIF = variance inflation factor, and COR = correlation matrix value.

Additionally, we reran all models on our three subsets of data: one that excluded species traditionally used in aquaculture, one that excluded those capable of air-breathing, and one that included only species for which gill surface area was measured across an order of magnitude of body size. Using these data subsets had no effect on our results (Tables S2 – S4).

All models described above for all three questions were fit in R using the Stan probabilistic programming language in *rstan* (Stan Development Team 2019; R Core Team 2020); see the SI for more detail on our modeling approach.

## Results

### 1. Do species with faster growth coefficients for their maximum body size have larger gills?

Gill surface area explained some variation in von Bertalanffy growth coefficient across species, regardless of which gill surface area metric was used: the ontogenetic intercept or the ontogenetic slope. While the 95% Bayesian Credible Intervals (BCI) of the gill surface area metric in each model overlapped with zero, a relatively small proportion of the posterior distributions of the effect sizes for the intercept and slope were greater than zero (intercept: 13%, slope: 25%; (Table 1, Figure 1). In contrast, asymptotic size explained substantial variation in growth coefficients across species in both models with none of the posterior distributions overlapping zero (Table 1, Figure 1). Based on the mean effect size estimates (slope values in Table 1), asymptotic size explained 3.25 times more variation than the ontogenetic intercept and 5.4 times more variation than the ontogenetic slope. Additionally, no multicollinearity or correlation was detected between gill surface area and asymptotic size in any of the three models (Table 1).

**Figure 1.**
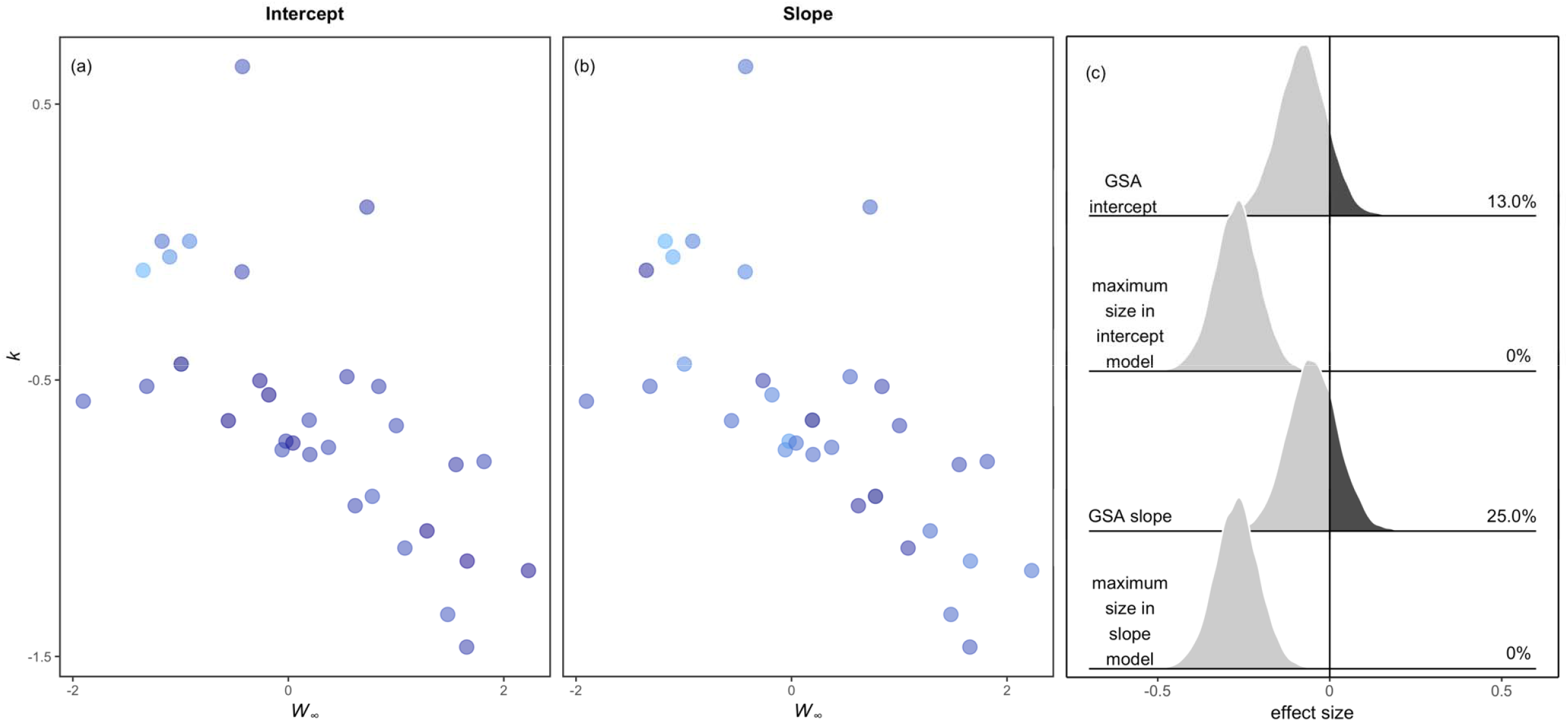
Gill surface area explains some variation in growth coefficients at a given maximum size. Gill surface area, as measured by (a) the species-specific ontogenetic intercept or the (b) ontogenetic slope of the relationship of gill surface area and body mass differs across fishes with different von Bertalanffy growth coefficients (*k*) and asymptotic sizes (). Gill surface area is indicated by a gradient of color, with darker blue indicating a greater intercept or slope, and lighter blue indicating a lower intercept or slope. Species-specific gill surface area intercepts and slopes and their relationships with *k* and were estimated in a Bayesian multilevel model where the first level estimated the ontogenetic intercept or slope and the second level estimated the relationship of *k*,, and the ontogenetic intercept or the ontogenetic slope. (c) The entire posterior distribution of each effect size, as well as the percent greater than zero (shaded dark grey) for both models in (a) and (b). Intercepts and slopes were standardized in the model prior to the second level (see text and SI for more detail).

### 2. Do estimates of gill surface area that incorporate its size-dependency (the slope and intercept of the ontogenetic allometry) relate to growth performance across fishes?

The relationship between gill surface area and growth performance was weakly positive for both metrics of gill surface area -- the ontogenetic intercept and the ontogenetic slope (Table 2, Figure 2). Although the 95% BCI of the effect size of gill surface area in both relationships crossed zero, the mean effect size for both was positive (and similar). Additionally, a large proportion of the posterior distributions for both gill surface area effect sizes were greater than zero. For the relationship of growth performance and the ontogenetic intercept, 93.5% of the posterior distribution was greater than zero (the mean slope = 0.36; 95% BCI -0.11 to 0.83, Table 2, Figure 2) and for the relationship of growth performance and the ontogenetic slope, 89.1% of the posterior distribution was greater than zero (the mean slope = 0.33 (95% BCI -0.20 to 0.83, Table 2, Figure 2).

**Figure 2.**
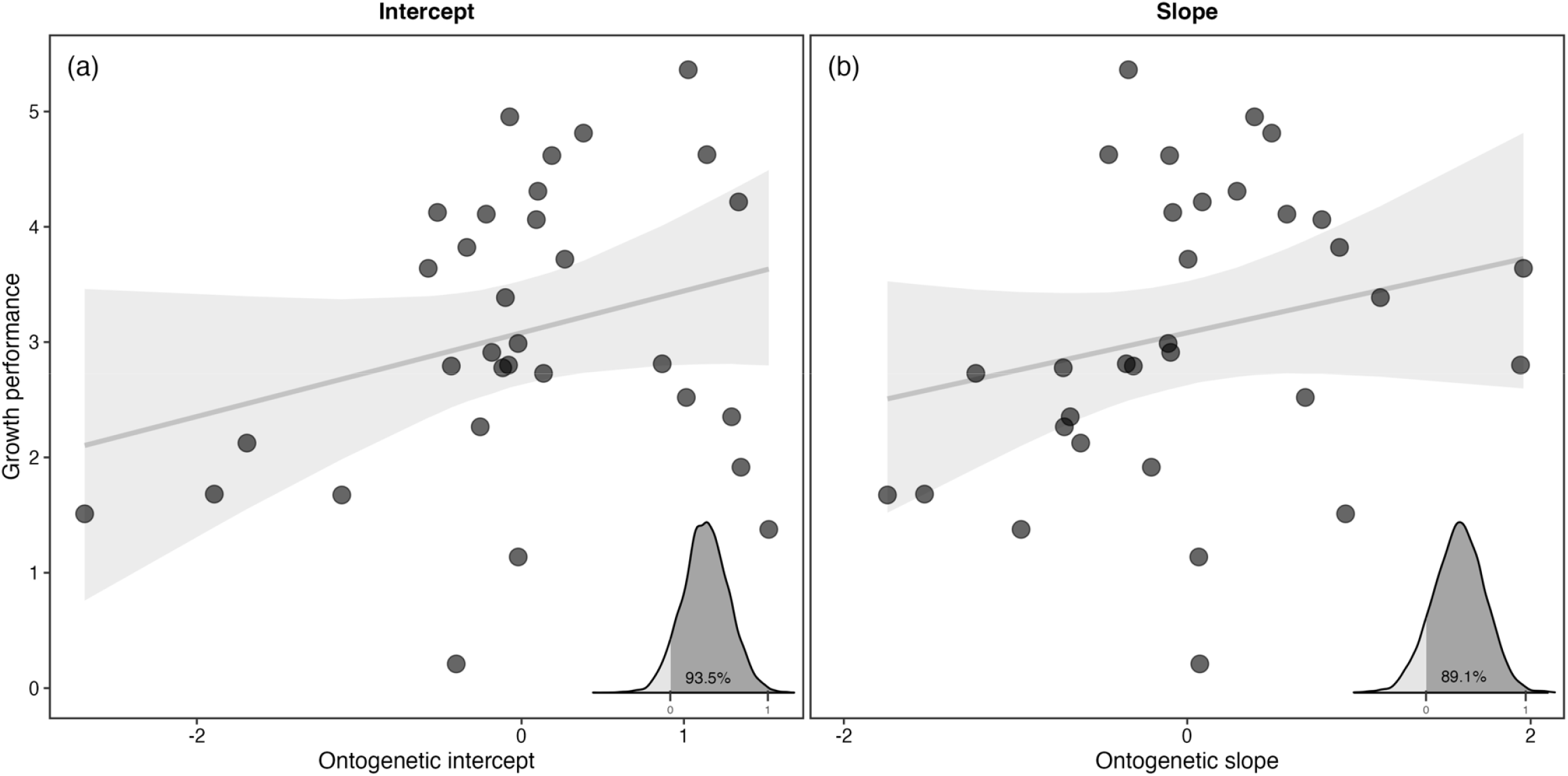
Gill surface area explains some variation in growth performance across fishes. The relationship between growth performance and (a) gill surface area ontogenetic intercept and (b) gill surface area ontogenetic slope. Species-specific intercepts and slopes and their relationships with growth performance were estimated in a Bayesian multilevel model where the first level estimated the gill surface area ontogenetic intercept and slope, and the second level estimated the relationship of growth performance and either the ontogenetic intercept or the ontogenetic slope. All predictors were standardized prior to the second level of the model. In all panels, the fit lines represent the fitted growth performance for each value of the respective gill surface area measure, and the grey shaded region represents the 95% Bayesian Confidence Interval (BCI). The 95% BCIs for all models overlapped with zero but a fairly large proportion of the posterior distribution was positive (see Table 2 and inset in each panel).

**Table 2.** Comparison of all coefficients and their 95% Bayesian Credible Intervals (BCI) estimated from models parameterized with growth performance as the response variable and the species-specific slope or intercept as the response variable (differentiated in the ‘Model parameterization’ column) with and without the inclusion of a phylogeny. A Bayesian multilevel modeling framework was used to estimate all parameters in Stan using the package *rstan* in R v4.0.2. All intercepts and slopes were standardized in the model prior to the second level (see text and SI).

| **Model parameterization** | **Intercept (95% BCI)** | **Slope (95% BCI)** | **Phylogenetic signal ($\lambda$)** | **Proportion of posterior distribution > 0** |
| --- | --- | --- | --- | --- |
| Growth performance ~ intercept | 3.08<br>(2.63 to 3.53) | 0.36<br>(-0.11 to 0.83) | -- | 0.94 |
| intercept | 3.08<br>(2.62 to 3.52) | 0.36<br>(-0.13 to 0.83) | 0.50<br>(0.03 to 0.98) | 0.93 |
| slope | 3.08<br>(2.63 to 3.53) | 0.33<br>(-0.20 to 0.83) | -- | 0.89 |
| slope | 3.08<br>(2.63 to 3.52) | 0.33<br>(-0.21 to 0.84) | 0.50<br>(0.03 to 0.98) | 0.89 |

### 3. Does activity level better characterize the variation in growth performance across species compared to gill surface area?

In both models, caudal fin aspect ratio explained more variation in growth performance across fishes compared to gill surface as measured by the ontogenetic intercept or the ontogenetic slope (Figure 3, Table 3). Based on the mean effect size estimates (slope values in Table 3) caudal fin aspect ratio explained 5.5 times more variation than the ontogenetic intercept and 3.3 times more variation than the ontogenetic slope. The 95% BCI of the effect sizes for the ontogenetic intercept and slope of gill surface area overlapped with zero yet a large proportion of the posterior distribution was greater than zero (68.3% and 79.6% for the ontogenetic intercept and ontogenetic slope, respectively; (Figure 3, Table 3). No multicollinearity or correlation was detected for any model that included both caudal fin aspect ratio and gill surface area based on variance inflation factor (VIF) or correlation indices (whether measured by the gill surface area ontogenetic intercept or the gill surface area ontogenetic slope; Table 3).

**Figure 3.**
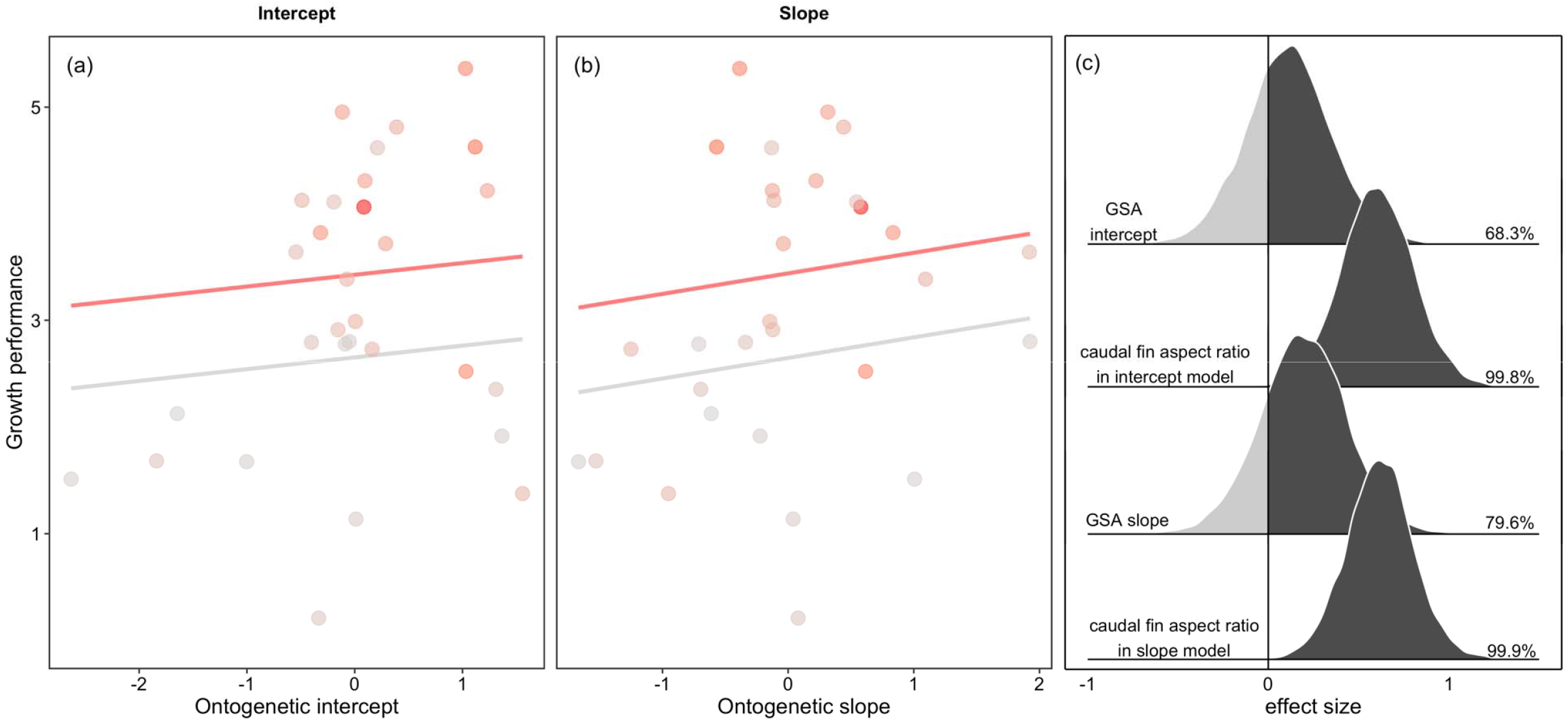
Activity level (as measured by caudal fin aspect ratio) explains more variance in growth performance compared to gill surface area, whether measured as the ontogenetic intercept or the ontogenetic slope. The relationship of growth performance and (a) the gill surface area ontogenetic intercept and (b) gill surface area ontogenetic slope, with each species’ respective value shaded according to their caudal fin aspect ratio value, with red indicating a high caudal fin aspect ratio (high activity level) and grey indicating a low caudal fin aspect ratio (low activity level). The fit lines in both plots correspond to the predicted growth performance at a given value of the respective gill surface area metric for the first (grey) and third (red) quartile of caudal fin aspect ratio. (c) The entire posterior distribution of each effect size, as well as the percent greater than zero (shaded in dark grey) for both models in (a) and (b). Gill surface area intercepts and slopes were standardized in the model prior to the second level (see text and SI for more detail).

**Table 3.** Comparison of coefficients and their 95% Bayesian Credible Intervals (BCI) for the relationship of growth performance, activity level (as measured by caudal fin aspect ratio), and gill surface area, as measured by the ontogenetic intercept or slope. All models were estimated using a Bayesian multilevel modeling framework in Stan using the package *rstan* in R v.4.0.2. All predictors in the second level of the model were standardized and thus the effect sizes for the slopes are relative to each other (see text and SI). GP = growth performance, CFAR = caudal fin aspect ratio, GSA = gill surface area, VIF = variance inflation factor, and COR = correlation matrix value.

| Model parameterization of the second level | Intercept (95% CI) | CFAR Slope (95% CI) | GSA slope (95% CI) | Phylogenetic signal ($\lambda$) | Relative importance of CFAR vs. gill surface area | VIF | COR |
| --- | --- | --- | --- | --- | --- | --- | --- |
| *growth performance ~ activity level +* |  |  |  |  |  |  |  |
| *intercept* | 2.95<br>(2.51 to 3.38) | 0.60<br>(0.21 to 1.00) | 0.11<br>(-0.35 to 0.58) | -- | 0.60/0.11 = 5.5 | 1.16 | 0.29 |
| *intercept* | 2.95<br>(2.52 to 3.39) | 0.60<br>(0.20 to 1.00) | 0.11<br>(-0.38 to 0.58) | 0.50<br>(0.03 to 0.98) | -- | -- | -- |
| *slope* | 2.95<br>(2.52 to 3.38) | 0.62<br>(0.25 to 0.99) | 0.19<br>(-0.29 to 0.67) | -- | 0.62/0.19 = 3.3 | 1.02 | 0.25 |
| *slope* | 2.95<br>(2.52 to 3.39) | 0.62<br>(0.24 to 0.99) | 0.19<br>(-0.29 to 0.65) | 0.50<br>(0.02 to 0.98) | -- | -- | -- |

## Discussion

Overall, we found that there was a positive relationship between gill surface area, growth, and maximum size across fishes; however, while gill surface area does explain some variation in growth performance, this relationship is weak. Other variables – particularly activity, were more important in explaining variation in growth and maximum size. While we used several models that progressively built on each other for each question, all models were in agreement and supported the idea that variation in growth and maximum size across fishes cannot simply be explained by gill surface area. We do note, however, that all models were quite noisy and had a great deal of residual error. This may provide further support that additional factors, such as those tested here (activity) or others (environmental temperature, food availability, reproduction), play a larger role in fish growth compared to gill surface area, and, that life history theory (the idea that life history traits trade off against one another in response to varying mortality and environments) remains valid, and may even be enough to explain variation in life history traits (Beverton & Holt 1959, Audzijonyte *et al*. 2019, Morais & Bellwood 2018, van Denderen *et al*. 2020). We next discuss what our results mean in the context of the Gill Oxygen Limitation Theory, what additional factors may be important in explaining variation in growth and maximum size, the pull between ecology and evolutionary history in shaping life history traits, the strengths of our modeling approach, and finally, end with suggested areas for future research.

From a life history perspective, our study suggests that gill surface area likely does not underlie maximum size in fishes, as predicted by the Gill Oxygen Limitation Theory. Instead, other mechanisms may underlie the suggested pattern of oxygen limitation on growth under warmer temperatures or in larger species (Hoefnagel & Verberk 2015; Audzijonyte *et al*. 2019, Rubalcaba *et al*. 2020). By extension, our work suggests that the scaling of gill surface area is unlikely to confer a limitation on the oxygen supply required for aerobic metabolism as it pertains to growth. However, the Gill Oxygen Limitation Theory is multifaceted (i.e., there are other parts to the theory such as ideas on protein denaturation, efficiency of assimilation, etc. that we did not test; Pauly 1981, 2010). Further, while we tested the central prediction of the Gill Oxygen Limitation Theory, there are other aspects of this theory – particularly those that have come to light as it has evolved since the 1980s – that remain to be evaluated (Pauly 2021). In addition, there remains much to be evaluated regarding oxygen limitation more broadly (i.e., not as it pertains to gill surface area), as well as the role of oxygen in shaping body size and other life history traits (Pauly 1981, 2010, 2021; Audzijonyte et al. 2019, Atkinson 1995). For example, Wong *et al*. (2021) found a weak link between resting metabolic rate (oxygen consumption) and growth performance and Bigman *et al*. (2018) found a relationship between gill surface area, maximum size, habitat type, and activity level. Clearly, the interrelationships among oxygen consumption and demand (metabolic rate, and gill surface area) activity, and growth are difficult to test directly, especially in the context of heterogeneity in environmental temperature.

We found that although evolutionary history did not improve our understanding of variation in growth and maximum size, activity level -- as measured by the aspect ratio of the caudal fin -- did. Taken together, these results suggest that ecology (activity) explains more variation in life history compared to evolutionary history (at least for growth and maximum size). It is no surprise that activity explains variation in life history as activity is intertwined with life history traits (e.g., body size and growth), habitat, and even gill surface area (Gray 1954, Hughes 1984, Wegner 2011, Bigman et al. 2018). It is non-trivial to partition variance between activity and gill surface area as they are related, which may be reflected in our results despite the low correlation found between the specific predictors used for gill surface area and growth performance used in our models. In terms of evolutionary history, it may be that relatedness truly does not explain remaining variation in growth and maximum size, which instead, may be entirely related to ecological and physiological processes that are the result of local adaptations and independent of shared ancestry. However, the lack of variance explained by the underlying phylogenetic structure in this relationship could also be due to the model of evolution implemented to account for underlying phylogenetic structure in datasets. Typically, phylogenetic comparative methods (including ours developed and employed here) rely on the Brownian motion model of trait evolution to model the expected variance and covariance between species (Felsenstein 1985, Freckleton 2009, Harmon 2018). This model of evolution assumes that traits evolve along the phylogenetic tree through a random-walk process (Harmon 2018). Thus, species that are more closely related have had less time to diverge and thus will have trait values that are more similar (i.e., they co-vary) compared to distantly related species, whose trait values have been randomly drifting for a longer period of time (Symonds & Blomberg 2014). Other, and perhaps better, models of trait evolution exist, yet implementing them in practice is nontrivial (Harmon 2018). However, rapid advancements in the implementation of more complex comparative methods are occurring, which will undoubtedly open the door to exploring trait evolution in a broader sense, to include employing other models of trait evolution (Pennell & Harmon 2013). To further explore the interplay between ecology and evolutionary history, future work examining other, possibly more refined measures of both activity level (e.g., swimming speed, aerobic scope), growth (we used von Bertalanffy growth coefficients), and other models of trait evolution may shed more light on the relationship of gill surface area, activity, and growth in the context of evolution.

A strength of this study and our general approach is the comparison of ontogenetic/intraspecific relationships across species. The central tenet of the Gill Oxygen Limitation Theory is that the ontogenetic scaling of gill surface area and body mass limits the supply of oxygen for growth as an organism increases in size, ultimately determining its maximum size (Pauly 1981, 2010, 2021). Thus, it is necessary to examine the relationship of growth and maximum size in the context of an allometry (or scaling), as opposed to using a metric such as gill area index or a mass-specific measure of gill surface area (Pauly 1981, 2010, 2021). Another important reason why an allometric scaling approach is necessary is evident when considering scale: the Gill Oxygen Limitation Theory is focused on how the scaling of gill surface area within species drives patterns across species, while other theories surrounding the role that oxygen plays in structuring life histories, population dynamics, and ecosystem functioning, among other processes, are largely centered on species-level mean data (i.e., the Metabolic Theory of Ecology; Brown *et al*. 2004). A combined approach that has the flexibility to incorporate raw and mean data, such as the modeling approach used here, will go a long way in helping us to understand how raw data, and the ontogenetic scaling relationships they confer, scale up to structure patterns across species, communities, and ecosystems. Indeed, understanding the role that oxygen plays in the ecology, physiology, and evolution of fishes will require an integrated approach that allows us to scale up individual-level physiological and ecological data to species- and ecosystem-level patterns.

To this end, we outline three areas for future research that would help us to understand the role that oxygen may play in structuring the growth, maximum size, and more broadly, the life histories of fishes. First, there is an underappreciated complexity in estimating accurate and reliable ontogenetic regression slopes. Ideally, species-specific raw data spanning the entire body size range would be used to estimate an ontogenetic slope, yet these data are rarely available. Estimating accurate slope values is central to testing whether the scaling of gill surface area (or other size-dependent traits such as metabolic rate) affect ecological, physiological, and evolutionary patterns across species. Here, we took care to identify the number of individuals of a given species that were required to produce a reliable and reasonable slope estimate, and only used data to estimate regression coefficients for species that had the minimum number of individuals. We urge other researchers to take a similar approach when estimating ontogenetic slope values. Future work could build off our simulations to identify the minimum proportion of a species’ size range needed to estimate a reliable and reasonable slope value. Second, future studies could examine other factors (e.g., temperature, food availability, metabolic rate) that may underlie life history traits and maximum size across species and assess whether and how oxygen interacts with these processes (Audzijonyte *et al*. 2018, Verberk *et al*. 2020, Wong *et al*. 2021). Indeed, we have not dealt with environmental temperature, a factor known to be important in explaining variation in growth across fishes (van Denderen *et al*. 2020). Future work could examine the relationships among gill surface area, environmental temperature, and growth to assess whether gill surface area varies with environmental temperature (as does metabolic rate), and whether this interaction may explain some variation in growth. An additional factor that may affect scaling relationships of metabolic rate and gill surface area, and thus size and growth, is reproduction. Indeed, recent work has identified that reproductive output may modulate the scaling of population growth rate (Denéchère et al., 2022, Barrowclift et al. *in review*). Perhaps reproductive output plays an outsized role in the relationship between gill surface area, metabolic rate, and growth (for discussion of this topic, see Marshall & White 2019). Third, without complementary experimental studies that can manipulate abiotic factors such as oxygen, temperature, and food availability, it is difficult to identify the mechanisms that confer the observed correlational patterns (Audzijonyte *et al*. 2018, Bigman *et al*. 2021). Marrying macroecological and experimental work will help us understand the role that oxygen plays in structuring growth and other life history characteristics, and more broadly, the ecology, physiology, and evolution of organisms (Audzijonyte *et al*. 2018). Such work is incredibly timely in light of the uncertainty regarding how the physiology and ecology of fishes will determine the response of species to continued global environmental change (Verberk *et al*. 2020, Lefevre *et al*. 2021).

## Acknowledgements

We would like to thank Daniel Pauly for the many enlightening discussions on this topic, as well as the members of the past and present Dulvy Lab for insightful comments over the years. We would also like to thank Erik Mahan for help with caudal fin aspect ratio measurements. This project was funded by the Natural Sciences and Engineering Research Council of Canada and the Canada Research Chairs Program. Author contributions: J.S.B. and N.K.D conceived of and designed the project and analysis. J.S.B. collected the data and performed all analyses and visualizations. All authors contributed to the interpretation of results. J.S.B. drafted the manuscript and supplementary information with input from all authors. N.K.D. supervised the project.

## Data and code availability

All data and code necessary to reproduce the results in this study will be archived in github upon acceptance. We place no restrictions on data or code availability.

