## Supplementary material for "Gills, growth, and activity across fishes": SI: 08142022_Bigman_etal_ComprativeMS_SI.docx

**Supplementary Information**

This file includes:

Supplementary Methods

Supplementary Figures: S1

Supplementary Tables: S1 – S4

**Supplementary Methods**

***Model Overview***

We constructed and compared Bayesian multilevel (i.e., hierarchical) linear regression models in R v.4.0.2 in Stan using the package *rstan* (Stan Development Team 2019; R Core Team 2020).

*i. Model Parameterization*

All multilevel models constructed in this study shared the same foundation and then were built upon to either add additional covariates (e.g., activity level) or a phylogenetic random effect. After the first level of the model, the relevant gill surface area metric was extracted (ontogenetic intercepts or ontogenetic slopes of gill surface area for each species) or calculated (gill area index). See text for a more detailed overview of how gill area index is calculated.

Basic parametrization

*First level of the model:*

*GSA_i_* = α + Σ*_j_*β*_j_x_i,j_* + ε_i_

$\hat{\varepsilon}$~ normal ($0, \sigma_{e}^{2}$)

α	~ student-t (3, 0, 10)

β*_mass_*	~ student-t (3, 0, 10)

$\sigma_{e}^{2}$ ~ half-Cauchy (0, 10)

Here, *GSA_i_* is the response variable (mean whole-organism gill surface area), α is the intercept, and β*_mass_* is the slope of the body mass associated with gill surface area, *x_mass_*. The choice priors for the intercept, α, slope, β*_mass_*, and error, $\sigma_{e}^{2}$, are explained below.

*Second level of the model:*

*Y_i_* = α + Σ*_j_*β*_j_x_i,j_* + ε_i_

$\hat{\varepsilon}$~ multivariate normal ($0, \sigma_{e}^{2}$)

α	~ student-t (3, 0, 10)

β*_j_*	~ student-t (3, 0, 10)

$\sigma_{e}^{2}$ ~ half-Cauchy (0, 10)

Here, *Y_i_* is the response variable (or growth performance, depending on the question), α is the intercept, and β*_j_* is the slope of the predictor (either the ontogenetic intercept or the ontogenetic slope of gill surface area, or, the gill area index for each species), *x_i,j_* for each species. The choice of priors for the intercept, α, slope, β*_j_*, and error, $\sigma_{e}^{2}$, are explained below. Other covariates (caudal fin aspect ratio and asymptotic weight) were added on as needed.

Models with phylogenetic parameterization

*Second level of the model:*

*Y_i_* = α + Σ*_j_*β*_j_x_i,j_* + ε_i_

$\hat{\varepsilon}$~ multivariate normal ($0, \sigma_{e}^{2}$)

α	~ student-t (3, 0, 10)

β*_j_*	~ student-t (3, 0, 10)

$\sigma_{e}^{2}$ ~ half-Cauchy (0, 10)

Here, the first level of the model does not change, but the second level does. Here, the second level of the model is as described above with the change in the residual error structure. Following (Frishkoff *et al.* 2017, Bigman *et al.* 2021), we assumed the residual error, ε_i_, to be distributed according to a multivariate normal distribution, where $\hat{0}$is a vector with length N, $\sigma_{e}^{2}$ is the variation in responses to the predictors (β_j_x_i,j_ ), and C_phylo_ is the N× N correlation matrix resulting from the phylogeny. The strength of the phylogenetic signal, λ, in the residuals under a model of evolution of Brownian motion is estimated according to C_phylo_ = λ * V + (1 - λ) * I, where V is the variance covariance matrix from the phylogeny, and I is an identify matrix of N × N values with $\sigma_{e}^{2}$ on the diagonal.

*ii. Choice of Priors*

We used weakly informative regularizing priors based on recommendations for Stan (<https://github.com/stan-dev/stan/wiki/Prior-Choice-Recommendations>). As λ (phylogenetic signal) has an equal chance of taking any value within the bounds of zero to one, we used a prior with a uniform distribution from zero to one. As $\sigma_{e}^{2}$ (variation in responses to the predictors (β*_j_x_i,j_*) can only be positive, we used a half-Cauchy prior with a location of zero and a scale of ten. Priors are also shown below for each set of models.

***Simulations***

To assess the number of individuals required to estimate a reliable slope estimate for the relationship between gill surface area and growth, we simulated how the estimated slope (and intercept) value varied with the number of individuals included in its estimation, based on a simple linear regression $(y= \beta_{0}+ \beta_{1}*x_{1}+ \varepsilon$).

To do so, we first defined a dataset of 100 individuals that ranged in body mass over four orders of magnitude (10g – 100000g). Next, we simulated the gill surface area data for these individuals at their given body mass based on simulated random errors and defined regression coefficients. For this, we simulated random errors for each unique observation (n = 100) from a normal distribution with a mean of zero and a standard deviation of 0.08. This value for the standard deviation was chosen as it was the mean standard deviation (sigma) for all gill surface area – body mass regressions for all species in our dataset with at least eight individuals (this threshold was chosen based on Jenkins & Quintana-Ascencio 2020). Similarly, we set the regression coefficients (intercept and slope) to the mean values estimated from these same regressions (mean slope = 0.85, mean intercept = 0.90). Next, we simulated the regression using the body mass and gill surface area data 1000 times and assessed the accuracy of the predicted regression coefficients (i.e., did the estimated regression coefficients match the defined regression coefficients?).

Second, we repeated these simulations 1000 times each on subsets of the dataset where a specified number of individuals (0 – 97) were dropped. In other words, we simulated a regression 1000 times on a random subset of three individuals (the minimum number to obtain a standard error on the regression coefficients), then four individuals, then five, etc., all the way up to 100 individuals. Finally, we plotted the standard error of the mean slope and intercept for each of these regressions (Figure S1).

Our simulations suggested that a threshold of eight individuals was sufficient for estimating reliable ontogenetic slope coefficients. We note, however, that this threshold of eight individuals, at least in our data simulations, is for a random spread of gill surface area and body mass (e.g., selecting individuals at random) as opposed to selecting a range of body size (e.g., individuals that span at least 33% of size range of species). Thus, a threshold of eight individuals of the same species to estimate an ontogenetic slope is likely conservative (i.e., fewer individuals may also result in a reliable slope based on the size range encompassed) yet produced reliable slope estimates that are within the known range of gill surface area slope values.

**Supplementary Figures**

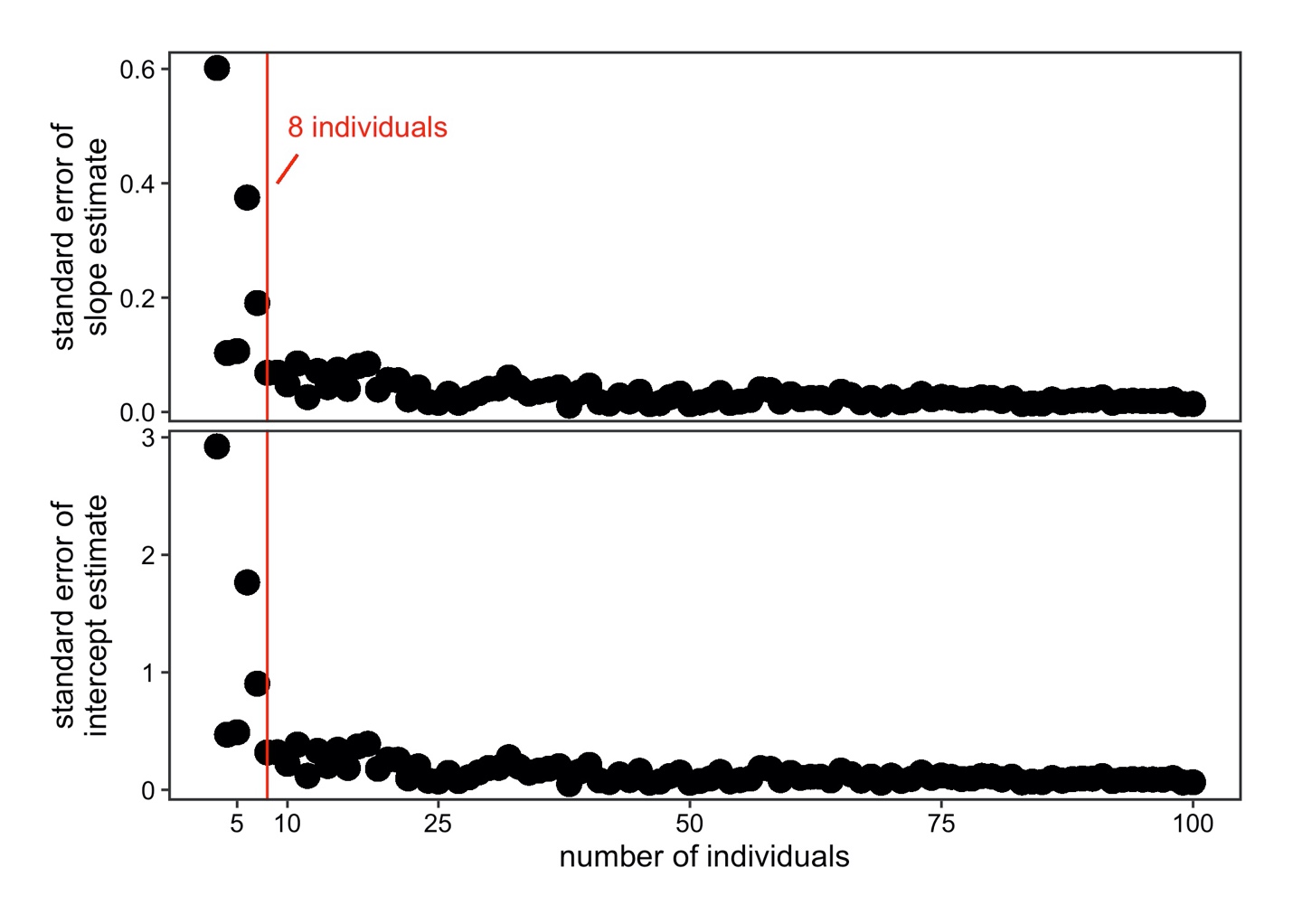

**Figure S1.** The change in the standard error of the slope (top row) and intercept (bottom row) resulting from simulations of the ontogenetic relationship between gill surface area and body mass with an increasing number of data points (individuals) included. The vertical red line on each plot indicates the threshold of eight individuals. For more detail, please see the Supplementary Methods.

**Supplementary Tables**

**Table S1**. The 32 species for which raw gill surface area and body mass data was available for at least eight individuals and for which von Bertalanffy growth parameters and phylogenetic position are known (see text). All models using this raw dataset were also compared with and without the inclusion of the species capable of air-breathing (and without those species traditionally used in aquaculture, which are indicated: *air-breather, ^+^aquaculture.

| **Scientific name** | **Common name** |
| --- | --- |
| *Acanthocybium solandri* | Wahoo |
| *Acipenser transmontanus* | White Sturgeon |
| *Alopias superciliosus* | Bigeye Thresher |
| *Alopias vulpinus* | Common Thresher |
| *Anabas testudineus** | Climbing Perch |
| *Anguilla anguilla** | European Eel |
| *Barbatula barbatula* | Stone Loach |
| *Boleophthalmus boddarti** | Boddart's Goggle-Eyed Goby |
| *Carcharhinus acronotus* | Blacknose Shark |
| *Carcharhinus isodon* | Finetooth Shark |
| *Carcharhinus limbatus* | Blacktip Shark |
| *Carcharhinus obscurus* | Dusky Shark |
| *Carcharhinus plumbeus* | Sandbar Shark |
| *Carcharodon carcharias* | White Shark |
| *Cirrhinus mrigala^+^* | Mrigal Carp |
| *Cobitis taenia** | Spined Loach |
| *Galeocerdo cuvier* | Tiger Shark |
| *Gymnocephalus cernua* | Ruffe |
| *Heteropneustes fossilis** | Stinging Catfish |
| *Hoplias malabaricus* | Trahira |
| *Isurus oxyrinchus* | Shortfin Mako |
| *Lipophrys pholis** | Shanny |
| *Micropterus dolomieu* | Smallmouth Bass |
| *Opsanus tau* | Oyster Toadfish |
| *Oreochromis niloticus^+^* | Nile Tilapia |
| *Petrocephalus catostoma* | Churchill |
| *Prionace glauca* | Blue Shark |
| *Rhizoprionodon terraenovae* | Atlantic Sharpnose Shark |
| *Scomber japonicus* | Pacific Chub Mackerel |
| *Sphyrna tiburo* | Bonnethead Shark |
| *Tinca tinca^+^* | Tench |
| *Torpedo marmorata* | Marbled Electric Ray |

**Table S2**. Comparison of coefficients and their 95% Bayesian Credible Interval (BCI) for the relationship of the growth coefficients (*k*), asymptotic size ($W_{\infty}$), and gill surface area, as measured by the ontogenetic intercept and the ontogenetic slope for models with the (a) full dataset, (b) full dataset excluding those species that are used in aquaculture, and the (c) full dataset excluding those species that are capable of air-breathing. All models were estimated using a Bayesian multilevel modeling framework in Stan using the package *rstan* in R v.4.0.2 All predictors in the second level of the model were standardized and thus the effect sizes for the slopes are relative to each other (see text and SI). GSA = gill surface area.

| **Model parameterization of the second level** | **Intercept**  **(95% BCI)** | $W_{\infty}$ **slope (95% BCI)** | **GSA slope (95% BCI)** | **Dataset** | **Sample size** |
| --- | --- | --- | --- | --- | --- |
| *k ~* $W_{\infty}$*+* |  |  |  |  |  |
| *intercept* | -0.55  (-0.68 to -0.42) | -0.26  (-0.38 to -0.13) | -0.08  (-0.22 to 0.06) | full | 30 |
| *intercept* | -0.62  (-0.71 to -0.52) | -0.26  (-0.35 to -0.17) | -0.06  (-0.17 to 0.04) | excluding aquaculture species | 27 |
| *intercept* | -0.50  (-0.67 to -0.33) | -0.34  (-0.50 to -0.18) | -0.07  (-0.22 to -0.08) | excluding air-breathing species | 25 |
| *slope* | -0.55  (-0.68 to -0.42) | -0.27  (-0.40 to -0.14) | -0.05  (-0.19 to 0.09) | full | 30 |
| *slope* | -0.61  (-0.71 to -0.52) | -0.27  (-0.36 to -0.18) | -0.01  (-0.12 to 0.11) | excluding aquaculture species | 27 |
| *slope* | -0.50  (-0.67 to -0.33) | -0.34  (-0.50 to -0.18) | -0.01  (-0.17 to 0.15) | excluding air-breathing species | 25 |

**Table S3.** Comparison of coefficients and their 95% Bayesian Credible Intervals (BCI) for the relationship of growth performance and the ontogenetic intercept and the ontogenetic slope estimated with the (a) full dataset, (b) full dataset excluding those species that are used in aquaculture, the (c) full dataset excluding those species that are capable of air-breathing, and (d) the dataset of raw species based on size range as opposed to number of individuals (see text). Both models were estimated using a Bayesian multilevel modeling framework in Stan using the package *rstan* in R v.4.0.2 All intercepts were standardized in the model prior to the second level (see text and SI).

| **Model parameterization of the second level** | **Intercept**  **(95% CI)** | **Slope (95% CI)** | **Dataset** | **Sample size** |
| --- | --- | --- | --- | --- |
| *GP ~* |  |  |  |  |
| *intercept* | 3.08  (2.63 to 3.53) | 0.36  (-0.11 to 0.83) | full | 32 |
| *intercept* | 3.06  (2.56 to 3.55) | 0.35  (-0.18 to 0.87) | excluding aquaculture species | 29 |
| *intercept* | 3.40  (2.94 to 3.87) | -0.05  (-0.56 to 0.46) | excluding air-breathing species | 26 |
| *intercept* | 3.22  (2.81 to 3.63) | 0.40  (-0.05 to 0.83) | species with a body size range of at least an order of magnitude | 34 |
| *slope* | 3.08  (2.63 to 3.53) | 0.33  (-0.20 to 0.83) | full | 32 |
| *slope* | 3.05  (2.56 to 3.54) | 0.32  (-0.27 to 0.88) | excluding aquaculture species | 29 |
| *slope* | 3.40  (2.94 to 3.84) | 0.23  (-0.33 to 0.78) | excluding air-breathing species | 26 |
| *slope* | 3.22  (2.79 to 3.65) | 0.20  (-0.29 to 0.67) | species with a body size range of at least an order of magnitude | 34 |

**Table S4**. Comparison of coefficients and their 95% Bayesian Credible Intervals (BCI) for the relationship of growth performance, caudal fin aspect ratio, and gill surface area, as measured by the ontogenetic intercept and the ontogenetic slope for models with the (a) full dataset, (b) full dataset excluding those species that are used in aquaculture, and the (c) full dataset excluding those species that are capable of air-breathing. All models were estimated using a Bayesian multilevel modeling framework in Stan using the package *rstan* in R v.4.0.2 All predictors in the second level of the model were standardized and thus the effect sizes for the slopes are relative to each other (see text and SI). GP = growth performance, CFAR = caudal fin aspect ratio, and GSA = gill surface area.

| **Model parameterization of the second level** | **Intercept**  **(95% CI)** | **CFAR Slope**  **(95% CI)** | **GSA metric slope (95% CI)** | **Dataset** | **Sample size** |
| --- | --- | --- | --- | --- | --- |
| *GP ~ CFAR +* |  |  |  |  |  |
| *intercept* | 2.95  (2.51 to 3.38) | 0.60  (0.21 to 1.00) | 0.11  (-0.35 to 0.58) | full | 30 |
| *intercept* | 2.88  (2.41 to 3.35) | 0.63  (0.22 to 1.05) | 0.11  (-0.41 to 0.62) | excluding aquaculture species | 27 |
| *intercept* | 3.19  (2.71 to 3.64) | 0.50  (0.13 to 0.89) | -0.15  (-0.62 to 0.33) | excluding air-breathing species | 25 |
| *slope* | 2.95  (2.52 to 3.38) | 0.62  (0.25 to 0.99) | 0.19  (-0.29 to 0.67) | full | 30 |
| *slope* | 2.87  (2.42 to 3.33) | 0.65  (0.28 to 1.02) | 0.21  (-0.30 to 0.70) | excluding aquaculture species | 27 |
| *slope* | 3.20  (2.75 to 3.64) | 0.48  (0.10 to 0.86) | 0.20  (-0.29 to 0.70) | excluding air-breathing species | 25 |
